# Toward passive acoustic occupancy surveys of rare red uakari monkeys (*Cacajao ucayalii*): using automatic signal recognition and cluster analysis to organize signals?

**DOI:** 10.1101/2025.02.10.637398

**Authors:** M. Bowler, I. Casinhas, D.Y. Moreno, G. Mori, F. Briceño, J. León, B Gacheva

## Abstract

We tested a workflow using a cluster analysis to develop a classifier to detect red uakari monkey calls using clustering and Hidden Markov Models, and Kaleidoscope viewer to review and discount false positives. We used recordings collected in a series of behavioral studies on a habituated group of red uakari monkeys on the Yavari-Miri River in the Peruvian Amazon as training data to develop the classifier and tested it on a passive acoustic survey at the same site where uakari distributions were known to us. We estimated detection probabilities for red uakaris and used an occupancy model to estimate habitat use to compare with use determined by behavioral research. We assessed a workflow for processing passive acoustic primate surveys that would eliminate false positives and demand minimal time and coding expertise from those implementing the analyses and reviewing audio recordings. These are key considerations in the design of landscape-scale PAM surveys for rare species.

## Introduction

Large mammal populations in tropical forests have been most commonly surveyed using direct observations within line transect methods, or arrays of ‘camera traps’ placed systematically through the forest (Zwerts, 2021). Line transects have the advantage of providing an established method for estimating population densities using the Distance method (Buckland et al. 2015) but require extensive effort and are ineffective for rare, nocturnal or cryptic species (Jeliazkov, 2022). Camera trapping overcomes limitations for species that are nocturnal or otherwise difficult to observe but, unless individual animals are readily identified, methods for estimating abundance and density have been slow to develop and verify (Wearn et al., 2022; but see Palencia et al. 2021). As a result, many surveyors have used ‘occupancy’ (Ψ), the proportion of sites occupied by a species, to compare and monitor animal populations, accounting for imperfect detection through occupancy modelling (Mackenzie et al. 2002).

A key limitation of camera trapping is the difficulty of including arboreal fauna. Arboreal communities make up a large proportion of animals in tropical forests and include species of conservation concern, such as many primates (Kays and Allison, 2001). Attempts to deploy camera traps in trees have most commonly addressed behavioral questions or taken inventories (e.g., Gregory et al. 2017; Whitworth et al. 2016; Moore et al. 2021), and while efforts to monitor populations through occupancy modelling with arboreal camera traps are increasingly successful (e.g., Bowler et al. 2017; Moore et al. 2020; Chen et al. 2021; Bersacola et al. 2022), these surveys require specialist climbing techniques, and are demanding in terms of time, effort and resources. More critically, detection probabilities are dependent on researchers identifying likely routes of travel through the canopy for target species, resulting in low, and variable detection probabilities and, therefore, a requirement for more camera stations. Furthermore, because the most likely routes of travel are often species-dependent, optimizing detection for one species or species group can compromise that for other species.

Passive acoustic monitoring (PAM) promises to improve surveys for the most vocal species. Remote acoustic recorders can be deployed systematically at a landscape scale and subsequently reviewed for animal calls to determine presence at sampling locations (Teixeira et al. 2024). Equipment can be cheaper, lighter and simpler than camera traps, and can monitor arboreal and terrestrial species simultaneously. Standardization is more achievable, since detection does not depend on finding specific routes through the canopy for arboreal species, or animal trails on the ground for terrestrial species.

The number and length of recordings typically collected in PAM generally preclude manual review, and automatic signal recognition through machine learning is inevitably necessary for reviewing passive surveys of a scale appropriate for population monitoring (Gibb et al. 2019). Algorithms are now well developed for some taxonomic groups in some geographic regions (e.g., birds, Zhong et al., 2021; bats, Paumen et al., 2022; frogs, Melo et al., 2021) and are widely adopted. Surveys for non-volent mammals using automatic detection are rarer, but a wide range of automatic signal detection techniques have been shown to work, including deep learning (e.g., Bergler et al. 2022; Batist et al. 2024), clustering (e.g., Clink and Klinck, 2021), Gaussian mixture models and support vector machine (e.g., Kalan 2015; 2016), hidden Markov models (e.g., Enari et al., 2019), and template matching (e.g., Zamboli et al. 2023). The utility of machine learning methods for detecting species, compared to camera trapping and direct observation, is species and context dependent, but passive acoustic surveys for Japanese macaques (*Macaca fuscata*) and sika deer (*Cervus nippon*) (Enari et al. 2019) and Chimpanzees (*Pan troglodytes*) (Crunchant et al. 2020) have been more efficient at detecting target species than camera traps, in part due to far larger detection areas around each station. The relative benefits of any survey method are highly dependent on the ecology and behavior of the species, and for acoustic methods, the communication of the target species clearly determines the suitability of PAM (Zwerts et al. 2021).

Like camera trapping, PAM can produce a detection history for defined sampling periods that emulates repeat sampling and can be used in occupancy modelling (Mackenzie et al. 2002). The potential and advantage of combining PAM with occupancy modeling for surveying primates have been long recognized (Kalan, 2015). However, the development of occupancy surveys using PAM has been slowed by the size of datasets required and the time-consuming process of verifying and eliminating false positives. While algorithms can now be readily developed for any vocal species, reviewing detections and eliminating false positives has held up many research and conservation projects. For example, unsupervised approaches, such as convolutional neural networks, BirdNET, affinity propagation clustering, Support Vector Machines, and Gaussian Mixture Models, provided high numbers of false positive occurrences when classifying vocalizations of indris (Ravaglia et al. 2023), Yucatán black howler monkeys (Wood et al. 2023), female gibbons (Clink et al., 2023), chimpanzees, Diana monkeys, red colobus and king colobus (Heinicke et al. 2015, Kalan et al. 2015). Occupancy modelling is robust to non-detection, but very sensitive to false positives, which produce biased occupancy estimates (Gu & Swihart 2004; Alldredge et al. 2008; McClintock et al. 2010; Miller et al. 2012; Clement, 2016). Indeed, a key assumption of the methodology is that false positives do not occur (Royle and Link, 2006). While there are models that explicitly account for false positives (e.g., Clare et al., 2021), where false positive rates are high, it is essential to manually validate detections to ensure that only true positives are included in analyses (Kalan et al. 2015), often requiring the review of tens of thousands of detections (e.g., in owls, Baroni et al. 2020)

The importance of minimizing false positives, and the high volume of false positives relative to target signals outputted by automated detection, has meant that the time taken to manually review files has become a key consideration in analyzing audio surveys. Applying efficient workflows to datasets containing terabytes of data, from which detection algorithms generate thousands of false positives, is a key challenge in surveying wildlife in tropical forests, especially where species are rare or patchily distributed. These issues are compounded in primates due to the rarity of target signals in many species. Much of the progress made in surveying primates with PAM and occupancy modeling has been made in species that routinely make long-range calls to communicate between widely separated groups or subgroups (e.g., howler monkeys, *Alouatta pigra*. Wood et al. 2023; Chimpanzees, *Pan troglodytes*, Crunchant 2020; spider monkeys, *Ateles geoffroyi*. Lawson et al. 2023), resulting in high detection probabilities relative to other primates. However, relative to many species of birds, insects and amphibians, even these primate calls are rare, and finding rare signals in very large, noisy datasets has proven to be a bottleneck for surveyors and has slowed the implementation of these methods.

The red uakari monkey (*Cacajao ucayalii*) is a primate with a patchy distribution on broad landscape scales in Western Amazonia (Silva et al. 2021; Ennes et al. 2024) and is absent from most localities within its broad geographical range (Bowler et al. 2009; Mayor et al. 2015). At many sites where it does occur, it is seen only rarely (Heymann and Aquino 2010). The species travels in large groups, with large home ranges that change seasonally (Bowler et al. 2009), but where groups are present, their large group sizes can make them the most abundant species within a local area for survey periods (Puertas and Bodmer 1993). The non-uniform distribution of the species means that abundance and density estimates made from line transects for the species are not always meaningful on a landscape scale, due to insufficient independent sampling points or lines, and poorly defined survey areas (Puertas and Bodmer 1993; Buckland 2010; Bowler et al. 2013; Heymann and Aquino, 2010; Mayor et al. 2015). Cutting and surveying sufficient numbers of transects (10-20 as a minimum, Buckland et al. 2010), and over sufficiently wide areas to account for patchily distributed species, is challenging and attempts at this are increasingly rare in the region, as researchers increasingly turn to camera traps to monitor hunted species (Zwertz et al. 2021).

Uakaris are vocal primates (Bowler and Bodmer 2009; Bezerra, 2010; León et al. 2024) and PAM shows promise in achieving occupancy surveys with coverage over larger geographical areas. The most common call for red uakari monkeys, the *hic* call (Fontaine 1981, synonymous to the *Ca-Ca-Ca-Ca*, in *Cacajao calvus*, Ayres 1986), is a short-range contact call given by all independent age-sex classes in a wide range of social situations; during travel, when fruiting trees are discovered, and after loud noises or other uakari calls (León et al. 2024). Uakaris moving in flexible groups of 30-80 individuals in the Yavari-Miri produced *hic* calls at a mean rate of 0.24 calls/individual/minute during daylight hours, making uakari monkeys one of the most vocal primate species (León et al., 2024).

We tested a workflow using a cluster analysis in user friendly Kalidoscope Pro software to develop a classifier to detect uakari *hic* call signals using clustering and Hidden Markov Models, and Kaleidoscope viewer to review and discount false positives. We used recordings collected in a series of behavioral studies on a habituated group of red uakari monkeys on the Yavari-Miri River in the Peruvian Amazon, as training data to develop the classifier, and tested it on a PAM survey at a site where uakari distributions were well known to us (Bowler et al., 2009; 2012; Bernárdez-Rodriguez et al. 2021). Our aims were to estimate detection probabilities for red uakaris using the aforementioned method, and to assess a workflow for processing PAM primate surveys that would be robust to false positives and demand minimal time and coding expertise from those implementing the analyses and reviewing audio recordings. These are key considerations in the design of landscape-scale PAM surveys for rare species.

## Methods

### Acoustic Surveys

Surveys were made on the Yavarí-Mirín River, Loreto, Peru, close to its mouth where it joins the Yavarí River on the border with Brazil (S04°27.5′ W071°45.9′). The area is covered by continuous forest, made up of three main forest types; non-flooding *terra firme* forest, white-water *várzea* forest that floods between November and May each year, and permanently waterlogged forest dominated by *Mauritia flexuosa* palms. The site has 13 species of primates, and the red uakari monkey (*Cacajao ucayalii*) is locally abundant (Bowler et al. 2013). The area was the site of continuous behavioral research on red uakaris between 2003 and 2005 (Bowler et al., 2009; 2012) and regular research activity through to the time of this survey (e.g., León et al., 2024; Bernárdez-Rodriguez et al. 2021), so home ranges and site occupancy for the red uakaris were well known.

Nineteen passive acoustic recorders (SM4, Wildlife Acoustics, Maynard, MA) were fixed to trees at approximately 150 cm from the ground, between 11^th^ and 17^th^ August 2018. Recorders were spaced systematically 1.3 to 2 km apart in two groups, without consideration of the historical home ranges or known occupancy of the monkeys. The first group (9 recorders) was to the east of the community of Carolina (two houses and a police post) in the Lago Preto Conservation Concession, and the second group (10 recorders) in the forest around the community of Esperanza (Figure 1). Fourteen of the recorders were set to record continuously, round the clock, while five recorders were set to record continuously between 05:30 hrs and 09:00 hrs to test gains in battery life and allow a cursory look at any effects on detection (not analyzed here). Although we know rates are higher in the morning and during travel (León et al. 2024), detection at a recording station depends on the interactions between rates of calling, the movements of the monkeys and background noise, so it was not clear how detection would vary during the day. We recorded audio files of 60 minutes duration at a sample rate of 16 kHz in the .wav format. A sample rate of 16 kHz was used, rather than the 48 kHz standard used in behavioral studies, to reduce file sizes, at the expense of sounds above 8 kHz in frequency - well above the frequencies of uakari calls (León et al. 2024). Recording continued until the batteries were exhausted, or when the recorders were collected between 19^th^ and 22^nd^ November 2018.

**Figure 1.**
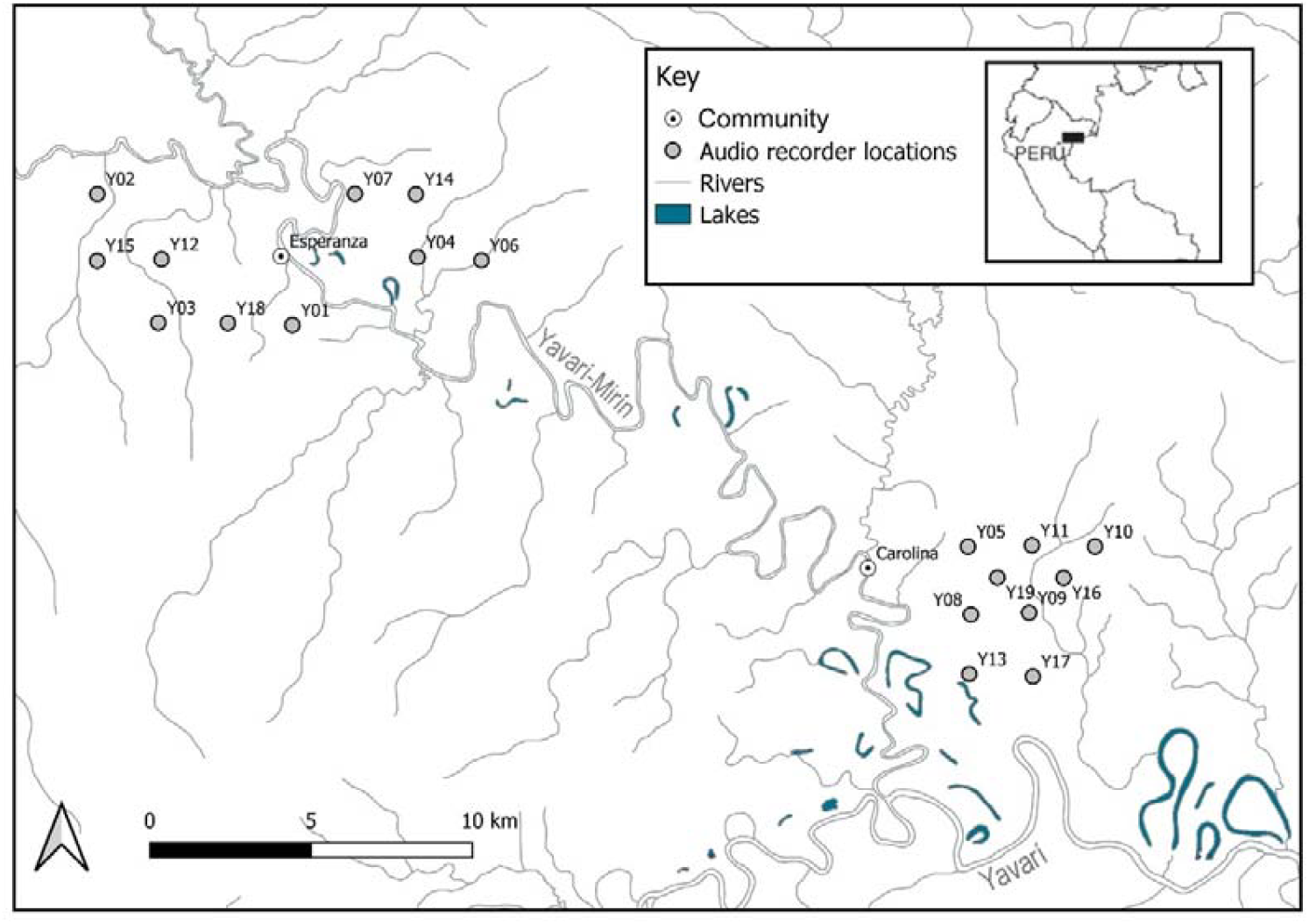
Remote audio recorder (SM4 Wildlife Acoustics) locations on the Yavarí and Yavarí-Mirín, Loreto, Peru.

### Audio Analysis in Kalidoscope

We used Kaleidoscope Pro vers. 5.4.6 (Wildlife Acoustics) to detect target signals in surveys and verify identifications to eliminate false positives. We selected Kalidoscope Pro because it offers a fast and efficient method for reviewing files. In analyses on acoustic surveys for American toads (*Anaxyrus americanus*), human time devoted to verifying positive identifications in Kaleidoscope Pro was around a tenth of that in a similarly user-friendly automatic identification software using BirdNET in Raven Pro (Pérez-Granados et al. 2023), which has been used successfully to analyze very small occupancy surveys to determine detection probabilities (Wood et al. 2023).

Kaleidoscope’s ‘cluster analysis’ groups the most common acoustic signals by similarity within customizable parameters. Because uakari calls are too rare, relative to other sounds in the forest, for Kaleidoscope Pro to cluster them from PAM recordings, we first created a ‘call classifier’ in Kaleidoscope by using the cluster analysis on audio recordings of red uakaris collected during behavioral research with habituated groups between 2003 and 2006 and in 2015 (León et al. 2024), in which uakari calls were among the most common signals.

We set parameters in Kaleidoscope to target the *hic* call (Figure 2). The *hic* call is somewhat variable in pitch but typically has around eight harmonics between 0.5 and 6 kHz. The second harmonic is often dominant at around 1.2-1.8 kHz and the 5^th^ is often the next loudest at around 3.3-4.2 kHz. The call is highly variable in the number of syllables (mean = 4.7 syllables per call, range 1-30 León et al. 2024). Syllables are 0.02-0.05 s in length, with an inter-syllable gap of 0.05-0.12 s.

**Figure 2.**
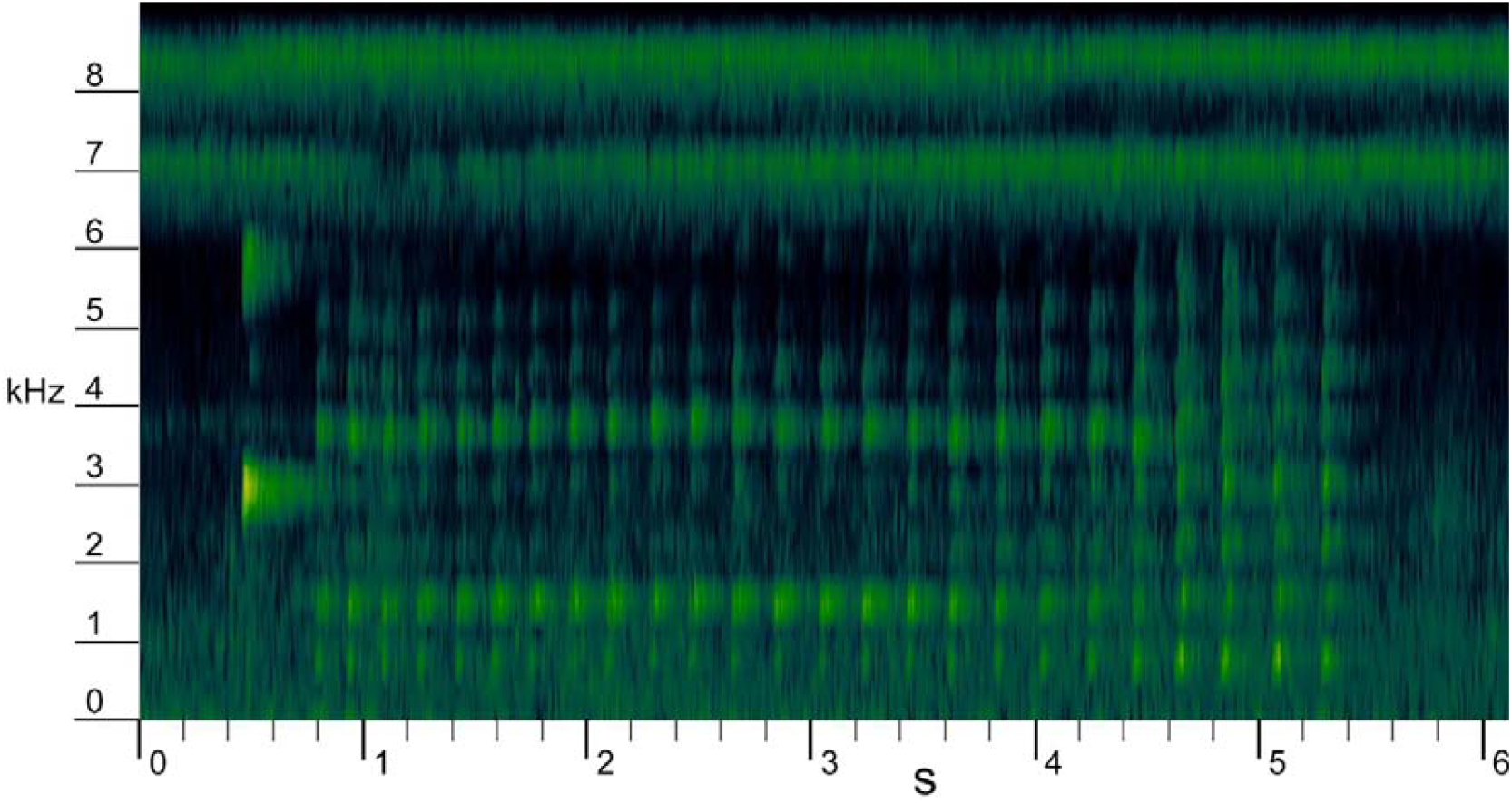
A *hic* call with 25 syllables given by a red uakari monkey (*Cacajao ucayalii*) during behaviouiral follows on the Yavarí-Mirín River, Loreto, Peru (S04°27.5′ W071°45.9′) from León et al. 2024. There are eight harmonics with primary and secondary harmonics at around 1.3-1.7 kHz and 3.5-4.0 kHz. This call is preceded by a *chick* call with a single syllable and a dominant harmonic at 2.8-3.2 kHz. *Chick* alarm calls typically elicit *hic* calls from other individuals.

**Figure 2.**
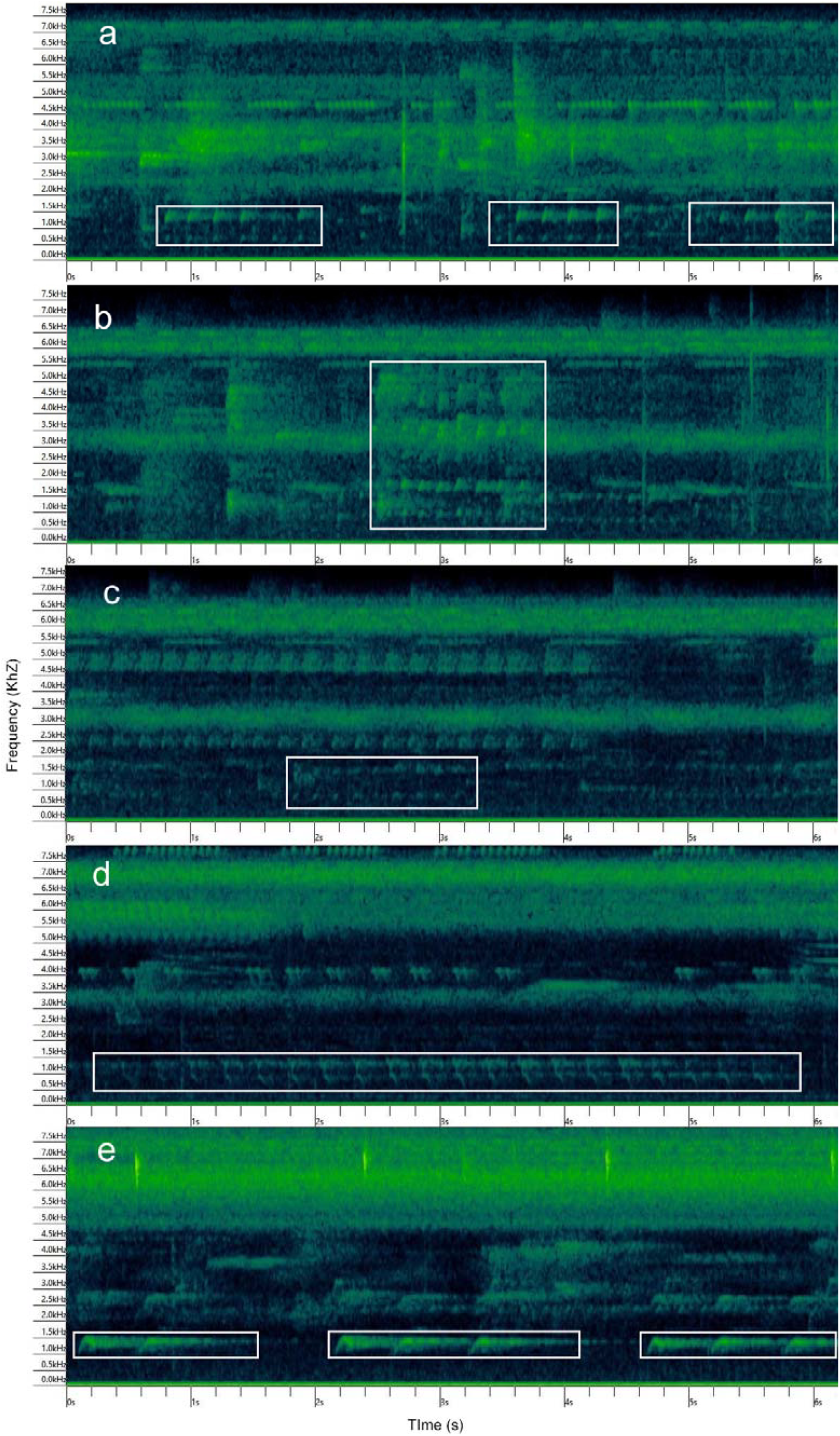
Examples of true positive and false positive detections of red uakari monkey (*Cacajao ucayalii*) *hic* calls returned by a call classifier in Kaleidoscope Pro vers. 5.4.6 (Wildlife Acoustics) from passive acoustic monitoring surveys on the Yavarí-Mirín River, Loreto, Peru (S04°27.5′ W071°45.9′), using SM4 audio recorders (Wildlife Acoustics) recording at 16 kHz. a-c true positive red uakari hic call detections of varying signal to noise ratios and showing different numbers of harmonics, d-e false positive detections of two birds commonly returned as red uakaris by our call classifier, blue-crowned Trogon*Trogon curucui* (d) and White-throated (Cuvier’s) Toucan *Ramphastos tucanus cuvieri* (e).

Because the *hic* call fills a broad range of frequencies, and was given in a noisy environment with frequent high-pitched signals from birds, frogs and insects, we targeted the lower two harmonics of the call by setting the minimum and maximum frequency range in Kaleidoscope to 0.5–2 kHz. We set a minimum and maximum length of detection to 0.5–15 s, and a maximum inter-syllable gap of 0.35 s.

We ran the cluster analyses on training data and reviewed clusters to manually identify and label call types. Kaleidoscope orders detections in order of ‘centrality’ to the cluster, so the first few recordings of each cluster indicate whether the cluster contained uakari *hic* calls. Clusters starting with other signals were ignored. The resulting classifier was applied to a further sample of training data and refined by manually identifying calls that Kaleidoscope included in these new *hic* clusters.

The refined call classifier was applied to PAM recordings from each recorder in turn. All 60-minute recordings from the same recorder were analyzed together in a single run of the classifier. Clusters that Kaleidoscope labelled as *hic* calls were manually verified visually in Kaleidoscope viewer by a single researcher (I. Casinhas). Those visually identified as likely uakari calls were played through Kaleidoscope and verified acoustically. Remaining true-positive detections were secondarily reviewed by another researcher (M. Bowler). Because Kaleidoscope orders call with the ‘most central’ to the cluster first, the user can quickly move between detections in order of the most likely true positives.

However, we did not limit our review of detections to the most likely detections. Although our occupancy model only required us to confirm presence for each recorder for each day, to compare numbers of detections in morning (05:30-09:30), midday (09:30-13:30) and afternoon (13:30-17:30) periods, and to assess the classifiers performance and rates of detection relative to false positives, we manually reviewed and labelled all detections labelled as uakaris by the classifier.

### Occupancy modelling

For each recorder, we combined detections for each day. So, for every day that a recording station was active, we had a detection history of 1 or 0. Because the low number of working sampling sites limited the effective use of covariates, we instead tested for differences in detection at different times of day using Chi Squared, before designing the model to include a measure of effort to account for the two different recording schedules used in the survey. Uakari activity starts shortly after 05:30 hrs and continues until around 17:30 hrs, at which point uakaris move very quietly to their sleeping trees (Bowler, 2007), so we considered recorders that were recording 24 hrs per day to have a diurnal recorder ‘effort’ of 12 hrs for our occupancy model, and recorders that were recording 05:30 hrs to 09:30 hrs to have an effort of 4 hours.

We ran a null single-season single species occupancy model in Presence Version 2.13.45 (USGS, 2023). Because our sample size (sampling points) was too low to include covariates such as habitat, we chose a null model in which ‘occupancy’ (Ψ) and detection (p) were uniform across sampling points. The aim was to estimate detection probabilities using our methods, for use in the design of future studies, and compare occupancy estimates with known patterns of presence and absence of red uakaris derived from behavioral research at the site (Bowler et al., 2009; 2012; León et al. 2024) and local ecological knowledge taken from interviews in the communities of Carolina and Esperanza (Bernárdez-Rodriguez et al. 2021).

## Results

Of the 19 recorders, three (all of which were set to record for 24 hrs per day) were damaged by water or termites and did not produce more than 48 hours of data, so were excluded from analysis.

Continuously recording stations (n=11) had a combined total of 230 recorder-days (5520 hours). The 4-hour per day recording stations (n=5) had 459 survey days (1836 hours).

Across all 7356 hours of recordings, our classifier in Kaleidoscope returned 100321 detections identified as uakari monkey *hic* calls. Manual review of all of these, taking approximately 80 hours, confirmed 356 uakari calls, while the rest were false positives (precision = 0.0035). True positive detections were present in 54 out of 7356 one-hour recordings, and on 39 recorder days.

Considering recordings from continuously-recording stations only, there was no statistical difference between the number of recordings with detections in the morning (n=7), midday (n=11) and afternoon (n=14) (*X2* (2, N = 32) = 2.3, *p* = 0.315). So we treated recorders only recording for 4 hours to have one third the ‘effort’ as recorders recording in all three periods. One detection occurred outside these diurnal periods, but at just one minute past the 17:30 cut-off.

Across the survey period, uakaris were detected at 11 out of 16 sites, with a naive occupancy of 0.688. Our null single season occupancy model estimated occupancy (Ψ) at 0.818 (standard error 0.131, confidence interval; 0.446-0.962), and a detection probability (*p*) of 0.068. Known uakari presence and absence at the site, inferred from intensive behavioral research and local ecological knowledge (Bowler et al., 2009; 2012; León et al. 2024; Bernárdez-Rodriguez et al. 2021), predicts uakari occurrence at 13/16 sampling points, while the remaining three points are isolated from uakari home ranges by the Yavari-Miri river, and absence is expected. This gives us an expected occupancy of 0.813 for the site. While the model estimate was very close to reality, it should be noted that the model confidence intervals were wide.

## Discussion

Our PAM survey produced an occupancy (Ψ) estimate for red uakaris (0.818) that very closely matched the known occupancy for the site (0.813). Confidence intervals were very wide, reflecting the low number of PAM recording stations that functioned (16/19). The precision of occupancy (Ψ) surveys using camera traps is highly sensitive to the level of species occupancy. Fewer than 20 camera sites may be sufficient for precise estimates of common (Ψ > 0.75) species, but more than 150 camera sites may be needed for rare (Ψ < 0.25) species (Kays et al. 2020). We expect PAM surveys to require similarly high numbers of recording stations.

Our temporal distribution of detections did not follow known patterns of calling recorded during behavioral follows, in which uakaris calls more in the morning (León et al, 2024) but may instead reflect periods when uakaris are more likely to move, and, therefore, be more likely to pass within range of a recorder. Alternatively, increased interference from other sounds during early morning may reduce detection probability. This small test provides no evidence for targeting periods with high call rates for this species and caution is advised before making this assumption for other species.

Manually verified positive uakari detections were present in only 54 out of 7356 one-hour recordings, which reflects the low density of groups of primates, even where a species is locally abundant relative to other primates, as with red uakaris at our site on the Yavari-Miri. While most recorders were in the known home ranges of uakari groups, uakari home ranges are large, at least 2200ha for one of the groups at our site (Bowler, 2012). Because uakaris are highly vocal when travelling, low detection probabilities at the site (0.068) likely reflect this wide ranging and infrequent visits to recording stations, rather than a low likelihood of detecting the monkeys when they were in range of the recorders. Other primates for which PAM occupancy surveys have been possible (e.g., howler monkeys, *Alouatta pigra*, Wood et al. 2023; Chimpanzees, *Pan troglodytes*, Crunchant 2020) have regular long-range calls, often given in the mornings, that are detected at long distances. This results in much higher detection probabilities and detection histories that are less dependent on the movements of the animals.

That uakari detection depends on their movements draws parallels with camera trapping surveys, and is significant, not only in determining detection probability, but also in how occupancy is interpreted and used to make inferences about populations. Occupancy models are difficult to fit for rare species, and for those with large home ranges, leading to overestimates for occupancy probabilities (Neilson et al. 2018, Stewart et al. 2018). In PAM surveys, as with camera trapping surveys, sampling units are typically undefined. Sampling units do not necessarily relate to the spacing of the detection devices, or the detection range of the device. As such, the assumption of geographic closure required by standard occupancy models (Mackenzie et al. 2002) is violated. It is not unusual for camera traps and passive audio recorders to be spaced at distances at which animals can be detected at several stations within a survey period, a fact utilized by analyses that use spatial capture-recapture methods (e.g., Green et al., 2020). Our recorder spacing at 1.3 to 2 km was within the maximum day ranges for red uakari monkeys (Bowler 2007). When the closure assumption is violated in this way, the probability of occupancy is re-interpreted as ‘probability of use’ (MacKenzie et al. 2002). However, this more precisely refers to the probability that the species was present at a sampling location at least once during the study period and is highly dependent on the length of the study period.

While designing shorter studies with more sampling points improves the utility of occupancy models for rare species (Ellis et al. 2014, Steenweg et al. 2018), this underutilizes the practical advantage of camera trap and PAM surveys in that they can be left for longer periods without additional cost (Emmet et al. 2021). It is likely that PAM surveys for species like the primates, which are often both rare and highly mobile, will require models that address both closure assumption violations and low detection probability simultaneously, such as continuous-time multi-scale occupancy models developed for low density carnivores (Emmet et al. 2021).

Uakaris occur across large remote regions of western Amazonia. The patchy distribution means that surveys will need wide coverage and large numbers of audio survey points. Implementing a workflow to process the large resulting datasets is a key step in achieving population assessments for this species and others with similar distributions. The necessity for extensive landscape scale surveys with large numbers of recording stations, and the requirement to minimize or eliminate false positives to obtain precise and unbiased occupancy estimates means that reducing false positives and facilitating manual verification of detections returned by any algorithm used is paramount. We demonstrated that it is possible to work through very large numbers of spectrograms visually in a reasonable timescale using Kaleidoscope viewer, and to produce occupancy estimates from a passive acoustic survey. However, as surveys increase in number and size, and estimates are required for large species assemblages, considerable manual verification time would be required until algorithms are further refined to reduce the large numbers of false positives that are currently typical. That said, it should not be necessary to review all detections. Because occupancy only requires ‘detection’ or ‘non detection’, manual review within a sampling period for a station can cease when a detection occurs. Although we reviewed all detections in this study, taking around 80 hours to review 100,321 spectrograms, clustering in Kaleidoscope orders the detections for centrality, enabling review of the most likely positive detections in order of confidence. Had we stopped on the first detection for each station during each sampling period (one day), the time take to review files manually would have been reduced to around thirty hours. It may additionally be acceptable to define a maximum number of files to review before recording non-detection, accepting a further reduction in detection probabilities. Other approaches include using citizen scientists to participate in the manual review, as used by online platforms for the annotation of large photo and video datasets for the detection of wild animals (Arandjelovic et al. 2016; Egna et al. 2020; Willi et al. 2019; Brookes et al. 2024).

As automatic signal detection techniques improve, more accurate detections will minimize false positives, but even low rates of false positives in surveys with many days of recordings will inevitably produce some false positives. Moreover, where practitioners use algorithms at new sites with different interference or variation in call structure, algorithm performance will vary, and without appropriate expertise in modifying algorithms, false positives will increase. Occupancy models are robust to lower and site-specific detection probabilities, but higher rates of false positives will require an efficient workflow to make acoustic surveys practicable for a wide range of end users in habitat countries.

These can clearly be developed, but for most end users, there is often a delay between published models and user-friendly workflows that researchers can implement to answer their ecological and conservation questions. We highlight the importance of a low cost, high efficiency, and wide usability of algorithms and technologies and workflows used for analyzing passive acoustic monitoring. We urge developers to consider the extra steps to scale up and disseminate their workflows for diverse users in habitat countries.

